# Reduction in olfactory ability in aging *Mitf* mutant mice without neurodegeneration

**DOI:** 10.1101/2024.06.24.600136

**Authors:** Fatich Mechmet, Eiríkur Steingrímsson, Pétur Henry Petersen

**Affiliations:** Department of Anatomy, BioMedical Center, Faculty of Medicine, University of Iceland, Reykjavik, Iceland; Department of Biochemistry and Molecular Biology, BioMedical Center, Faculty of Medicine, University of Iceland, Reykjavik, Iceland

**Keywords:** Age-related decline, olfactory bulb, olfactory function, *Mitf* mutation, neuronal hyperactivity, potassium channels

## Abstract

Age-related decline occurs in most brain structures and sensory systems. An illustrative case is olfaction, where the olfactory bulb (OB) undergoes deterioration with age, resulting in reduced olfactory ability. Decline in olfaction is also associated with early symptoms of neurodegenerative diseases including Alzheimer’s disease (AD) and Parkinson’s disease (PD). However, the underlying reasons are unclear. The microphthalmia-associated transcription factor (MITF) is expressed in the projection neurons (PNs) of the OB – the mitral and tufted (M/T) cells. Primary M/T cells from *Mitf* mutant mice show hyperactivity, potentially attributed to reduced expression of a key potassium channel subunit, *Kcnd3*/Kv4.3. This influences intrinsic plasticity, an essential mechanism involving the non-synaptic regulation of neuronal activity. As neuronal hyperactivity often precedes neurodegenerative conditions, the current study aimed to determine whether the absence of *Mitf* has degenerative effects during aging. Aged *Mitf* mutant mice showed reduced olfactory ability without inflammation. However, an increase in the expression of potassium channel subunit genes in the OB suggests that during aging compensatory mechanisms lead to stabilization.

**Significance statement:** This study highlights the age-related decline in olfaction and elucidates compensatory mechanisms mediated by potassium channels. These findings improve our comprehension of the processes underlying age-related changes in olfaction.

## Introduction

Aging is characterized by progressive deterioration. This includes loss of cognitive functions and reduction in the efficiency of sensory systems. Loss of sensory ability can also be a part of neuronal pathologies. Olfaction, the detection and discrimination of odors, shows an age-dependent decline in humans (Rawson et al., 2012; Schubert et al., 2017) and rodents (Patel & Larson, 2009), but olfactory impairment is also associated with a wide range of neurodegenerative and cognitive diseases, including Alzheimeŕs disease (AD) (Murphy, 2019; Tzeng et al., 2021) and Parkinsońs disease (PD) (Doty, 2012; Fullard et al., 2017). Moreover, partial or total loss of olfactory ability (hyposmia or anosmia) has been shown to contribute to depression (Sabiniewicz et al., 2022) and to affect daily life (Schäfer et al., 2021). Olfactory dysfunction is therefore both a possible biomarker of disease and a causal factor. It is therefore important to investigate underlying pathological mechanisms of olfactory dysfunction during aging.

The olfactory bulb (OB) is the first structure in the central nervous system (CNS) responsible for processing olfactory information. It is a well-defined and multi-layered structure, with well-characterized neuronal subtypes (Nagayama et al., 2014). The olfactory process starts at the olfactory sensory neurons (OSNs) in the olfactory epithelium (OE), which detect odor molecules and synapse with two kinds of projection neurons (PNs), the mitral and tufted (M/T) cells, as well as interneurons and periglomerular cells (PGCs) in the OB (Lodovichi & Belluscio, 2012). The M/T cells transmit information to the piriform cortex (PCx) and other olfactory associated brain areas, such as the olfactory tubercle (OT), amygdala, and orbitofrontal cortex (OCx) (Mori et al., 2013). Any changes in neuronal turnover, function or morphology can disrupt the OB circuitry triggering impaired olfaction. The known impairments which cause age-related decline in olfactory system include structural, molecular, and functional modulations in the OE, main olfactory bulb (MOB), and other regions that are involved in olfactory processing (Doty & Kamath, 2014).

The microphthalmia-associated transcription factor (*Mitf)* encodes a member of the *Myc* supergene family of basic helix-loop-helix zipper (bHLH-Zip) transcription factors (Goding & Arnheiter, 2019; Steingrímsson et al., 2004). The MITF protein, which is known as a master regulator of melanocytes, also plays central roles in mast cells (Oppezzo & Rosselli, 2021) osteoclasts (Lu et al., 2010; Steingrimsson et al., 2002), and has a distinct expression in the M/T cells (Atacho et al., 2020; Ohba et al., 2015). Previous studies have shown that *Mitf* plays a role in intrinsic plasticity of primary M/T cells of the mouse OB. Its loss leads to increased intrinsic excitability (Atacho et al., 2020). This suggests a role in homeostatic plasticity, which refers to the adjustment of the neuronal activity while keeping the activity of an individual neuron stable (Turrigiano, 2012). Primary PNs from *Mitf^mi-vga9/mi-vga9^* mice are hyperactive– likely due to a reduced expression of potassium channel subunits (Atacho et al., 2020). As neuronal hyperactivity is a hallmark feature of the early phase of neurodegeneration (Chase & Markopoulou, 2020; Hector & Brouillette, 2020; Marin et al., 2018; Targa Dias Anastacio et al., 2022), it is of interest to determine whether the lack of *Mitf* has detrimental effects in aged mouse OB. This question was addressed by examining markers of inflammation, evaluating the number of PNs and analyzing global gene expression. The results suggest a buffering of neuronal activity via compensatory mechanisms in aging mice. The study contributes the understanding behind the mechanism of aging in the olfactory system and the adaptation of CNS adaptation to dysregulated neuronal activity.

## Material and methods

### Animals

All *in vivo* procedures were approved by the Committee on Experimental Animals and were in accordance with Regulation 460/2017 and European Union Directive 2010/63 (license number 2013-03-01). Wild type (C57BL/6J), and homo- and heterozygous *Mitf^mi-vga9^* mutant mice were used in this study. Mice were housed at the mouse facility at the University of Iceland, in groups of two to three per cage under controlled conditions (21-22°C; 12 hours light/ 12 hours dark). Unless indicated otherwise, food and water were provided *ad libitum*.

### Behavior: Hidden food assay

Hidden food assay was performed during the active phase of mice in the dark cycle. Mice were first habituated to test environment by placing them individually in cages for 24 hours. Food and water were *ad libitum,* and Cocoa Puffs (brand name Nesquik General Mills) were placed in each cage in a small petri dish (4 pieces per cage) for 12 hours. The consumption of cereals was monitored for two consecutive days to ensure the cereals were palatable. Following overnight (O/N) starvation (experimental license number 2016-05-01), mice were kept in an odor-free room for 1 hour with no water, food, or cereals. Each mouse was tested separately. In the assay, a cereal was hidden in an opposite corner under the bedding in a new cage, and the time spent finding it was measured.

### Immunofluorescence (IF)

Mice were transcardially perfused (license number: 2014-07-02) with 1x phosphate-buffered saline (PBS; Gibco, cat# 18912-014) followed by 4% paraformaldehyde (4% PFA; Sigma-Aldrich, cat# P6148) in 1x PBS pH 7.4. Subsequently, brains were post fixed with 4% PFA in 1x PBS at 4°C O/N. Following this, the brains were rinsed with 1x PBS for two days. OBs from young and aged C57BL/6J and *Mitf^mi-vga9/mi-vga9^* mice were dissected and placed in a Peel-A-Way Disposable Plastic Tissue Embedding Molds (Polysciences, Inc.) filled with 5% agarose (Invitrogen, cat# 15517-014) dissolved in dH_2_O. The OBs were sectioned into 50 μm thin sections at room temperature (RT) using a microtome, Microm HM 650V (Thermo Scientific), and were kept at 4°C until further use. Sections were blocked in blocking buffer composed of 10% normal goat serum (NGS; Gibco, cat# 16210-064) and 0.3% Triton X-100 (Sigma, cat# T8787) in 1x PBS for 1 hour at RT and were then incubated with primary antibodies diluted in the blocking buffer at 4°C O/N. The primary antibodies used in this study were as follows: Rabbit polyclonal anti-GFAP (1:1000; abcam, ab2760), rabbit monoclonal [EPR16588] anti-Iba-1 (1:1000; abcam, ab178846), rabbit monoclonal [EPR21950] anti-Tbr2/Eomes (1:1000; ab216870), rabbit monoclonal [EPR8143(2)] anti-Tbr1 (1:1000; ab183032). Following three washing steps with 1x PBS for 5 minutes each, the sections were incubated with secondary antibodies, Alexa Flour 546 IgG anti-rabbit for anti-GFAP and anti-Iba-1, Alexa Flour goat goat 647 IgG anti-rabbit for anti-Tbr2/Eomes and anti-Tbr1 (Life Technologies; each diluted at 1:1000 in the blocking buffer), and DAPI (1:1000; Sigma, cat# D9542) for 1 hour at RT in the dark. After the wash with 1x PBS, the tissues were put onto Menzel-Gläser Superfrost Plus microscope slides (Thermo Scientific, cat# 2621573) and mounted using Fluoromount Aqueous Mounting Medium (Sigma, cat# F4680). Imaging was performed at 20x magnification using 20 confocal Z-stacks. Four images, two from lateral and two from medial OB, were obtained from each section.

### Golgi-Cox staining for light microscopy

Golgi-Cox staining was performed according to the protocol described by Vints and associates (Vints et al., 2019). Briefly, PFA-fixed OB sections from aged C57BL/6J and *Mitf^mi-vga9/mi-vga9^* mice were incubated in Golgi-Cox solution, a mixture of 5% mercury chloride (Sigma-Aldrich, cat# M1136), 5% potassium dichromate (Sigma-Aldrich, cat# 207802) and 5% potassium chromate (Sigma-Aldrich, cat# 216615) for fifteen days at RT in the dark. On the fifteenth day, the sections were washed three times for 5 minutes with dH_2_O and transferred to 28% ammonium hydroxide (Sigma-Aldrich, cat# 221228) in dH_2_O for 30 minutes at RT with rotation. Following three washing steps with dH_2_O, the sections were incubated in 15% ILFORD RAPID FIXER (Harman Technology Limited) in dH_2_O for 10 minutes at RT and then washed three times with dH_2_O. The sections were put onto Menzel-Gläser Superfrost Plus slides (Thermo Scientific, cat# 2621573), dried at RT, and mounted with Mowiol 4-88 (Aldrich, cat# 81381) mounting medium. Slides were imaged using a Leica light microscope.

### Western blotting

Mice were sacrificed by cervical dislocation. OBs from young and aged C57BL/6J and *Mitf^mi-vga9/mi-vga9^* mice were weighed and homogenized in radioimmunoprecipitation assay (RIPA) buffer (250 mM NaCl, 1% IGEPAL CA-630, 0.5% sodium deoxycholate, 0.1% sodium dodecyl sulphate, 50 mM Tris-HCl pH 8.0) containing 1x Halt protease & phosphatase inhibitor cocktail (Thermo Scientific, cat# 78440). Lysates were spun at 16,000 x g for 10 minutes at 4°C and the supernatant transferred to a new tube. After determining the protein concentration through a Bradford protein assay using Bradford reagent (Sigma, cat# B6916), each sample was diluted with an appropriate volume of RIPA buffer mixed with 1x Halt protease & phosphatase inhibitor cocktail to achieve a final protein amount of 20 µg protein. Following this, 2x sample buffer (4% sodium dodecyl sulphate, 20% glycerol, bromophenol blue 0.02%, 120 mM Tris-HCl pH 6.8) containing 5% β-mercaptoethanol (Aldrich, cat# M6250) was added to each sample at 1:1 ratio and the lysates were heated at 95°C for 5 minutes. The protein lysates were run on 12.5% SDS-PAGE gel and were transferred onto 0.2 µm PVDF membrane (Thermo Scientific, cat# 88520). The membrane was incubated in blocking buffer consisting of 5% non-fat dried milk in 1x Tris-buffered saline/0.1% Tween 20 (TBS-T) for 1 hour at RT. Following this, the membranes were incubated O/N at 4°C with the following primary antibodies diluted in the blocking solution: rabbit polyclonal anti-GFAP (1:1000; abcam, cat# ab7260), rabbit monoclonal [EPR16588] anti-Iba-1 (1:1000; abcam, ab178846 [EPR16588]), mouse monoclonal anti-beta Actin (1:5000; ab8224), mouse anti-GAPDH (1:5000; ab8245). The membranes were washed with 1x TBS-T and incubated for 1 hour at RT with the following secondary antibodies diluted at 1:10000 in the blocking buffer: IRDye goat anti-rabbit IgG 800 (green; LI-COR Biosciences), IRDye donkey anti-mouse IgG1 680 (red; LI-COR Biosciences). After washing steps with 1x TBS-T, the membranes were imaged using Odyssey CLx Imager (LI-COR Biosciences).

### RNA sequencing (RNA-seq) and data analysis

OBs from aged C57BL/6J and aged *Mitf^mi-vga9/mi-vga9^* mice were dissected, and flash frozen in liquid nitrogen. Tissues were homogenized in 1x DNA/RNA Protection reagent (New England Biolabs Inc., Part: T2011-1) and RNA was isolated using a commercially available kit (Monarch Total RNA Miniprep Kit; New England Biolabs Inc., cat# T2010S).

The quality (RIN score) and quantity of the isolated total RNA samples were evaluated using the DNA 5K/RNA chip on the LabChip GX instrument. To create cDNA libraries from Poly-A mRNA, Illumina’s TruSeq RNA v2 Sample Prep Kit (Illumina, RS-122-2001) was used. This process involved isolating Poly-A mRNA from the total RNA samples (0.2–1 μg input) through hybridization with Poly-T beads.

Subsequently, the Poly-A mRNA was fragmented at 94°C with divalent cations, followed by initiating first-strand cDNA synthesis using random hexamers and SuperScript IV reverse transcriptase (Invitrogen, cat# 18090010). Following this, second-strand cDNA synthesis, unique dual-indexed adaptor ligation, and PCR amplification were performed. The resultant cDNA sequencing libraries were evaluated on the LabChip GX, then diluted to 3 nM and stored at −20°C.

For the sequencing, the samples were pooled, and clustering carried out on NovaSeq S4 flow cells. The sequencing approach involved paired-end sequencing using the XP workflow on NovaSeq 6000 instruments (Illumina). Basecalling was performed in real-time using RTA v3.4.4. The process of demultiplexing BCL files and generating FASTQ files was done using bcl2fastq2 v.2.20.

A companion package of Kallisto (v 0.46.1) (Bray et al., 2016) was used to quantify transcript abundance following the pseudoalignment of RNA-seq data to the mouse reference genome (*Mus musculus*. GRCm38.96) (Yates et al., 2016). Differentially expressed (DE) genes in the OBs of aged C57BL/6J and *Mitf^mi-vga9/mi-vga9^* (n=3 per genotype) were identified using Sleuth (Pimentel et al., 2017). The significance of DE genes (p-/q- values) and the fold change (beta estimate) were determined using likelihood test (LRT) intersected in Sleuth.

Volcano plots were used to represent DE genes between aged *Mitf^mi-vga9/mi-vga9^*and C57BL/6J OBs by plotting the significance on the y-axis and cut-off of fold change on the x-axis (q-value ≤ 0.05; cut-off of |b-value (foldchange)|≥ 0.7) using R packages *“ggplot2”* and *“ggrepel”*. R package *“cowplot”* was used to create bar graphs showing TPM values of each gene between aged *Mitf^mi-vga9/mi-vga9^*and C57BL/6J. Functional enrichment analyses (GO terms) in molecular functions (MF) and enrichment analysis of pathways (KEGG) were generated combining p-value ≤ 0.05 and cut-off of |b-value (foldchange)|≥0.7 with *“clusterProfiler”* in the Bioconductor R package (Yu et al., 2012).

### Quantification of Tbr2/Eomes and Tbr1 positive cells

Tbr2/Eomes and Tbr1 positive cells from OB sections of young and aged C57BL/6J and *Mitf^mi-vga9/mi-vga9^* mice were counted using ImageJ software. Briefly, the software was uploaded with oib images of the lateral and medial OB, showing the GL, EPL, and MCL. After stacking the image at *“Maximum Intensity”* using *“Z-project”* analyzing method of the software, the image channels were split. Subsequently, a channel which shows Tbr2/Eomes or Tbr1 antibody staining was chosen. To make weaker stained cells more distinguishable, the channel was converted to grayscale and Tbr2/Eomes and Tbr1 positive cells in the GL, EPL and MCL were counted manually in the lateral and medial OB. The average number of cells per image was calculated.

### Analysis of Golgi-Cox images

Dendrite segmentation was performed using a pix2pix conditional generative adversarial network (cGAN) machine learning model with a resolution of 256×256 pixels and three-color channels. The training dataset included annotated sections from four images with random patches and augmentations applied during training. Model weights were saved every 50 steps to mitigate memory loss, and the best model was selected post-training. During inference, the model was applied multiple times across the image with different offsets to avoid artifacts. Segmentation accuracy was assessed qualitatively by experts on 32 images. Post-inference processing to quantify dendrite properties was performed using custom Python code and relevant libraries. These metrics were then analyzed for correlations with aged *Mitf^mi-vga9/mi-vga9^* compared to C57BL/6J.

### Statistical analysis

At least three mice per genotype and age group were included in the study. Quantitative results were analyzed using two-way ANOVA, and two-tailed unpaired Student’s t-tests using the R statistical package. To obtain p-values for ANOVA tests, multiple comparisons were conducted with Sidak’s and Tukey’s corrections. Numerical results represent the mean and the standard deviation (SD). The criteria for significance levels used to generate volcano plots and bar graphs from RNA-seq analysis are described both in the appropriate section of the Materials and Methods and in the corresponding figure legends. Sample size was not pre-determined prior statistical methods.

## Results

### Reduced olfactory ability is frequently observed in aged *Mitf^mi-vga9/mi-vga9^*

To assess the role of aging on the olfactory ability of *Mitf^mi-vga9/mi-vga9^*, a hidden food assay was performed with young and aged C57BL/6J, *Mitf^mi-vga9/+^*, and *Mitf ^mi-vga9/mi-vga9^*. Mice were divided into three age groups: 1-2, 8-12 and 14-27 months old. Diagrammatic representation of the assay setup can be seen in Figure 1A. Young *Mitf^mi-vga9/mi-vga9^* behaved similar to young C57BL/6J and *Mitf^mi-vga9/+^* mice whereas aged *Mitf^mi-vga9/mi-vga9^* often had significantly reduced olfactory ability compared to aged C57BL/6J. The olfactory ability of 8-12 months old *Mitf^mi-vga9/mi-vga9^* were lower than C57BL/6J by 15% (8-12 months old: *t _(80)_ = −2.584*, *p = 0.030*8). In the age range of 14-27 months, *Mitf^mi-vga9/mi-vga9^* mice showed 34.5% decrease in olfactory ability compared to C57BL/6J (14-27 months old: *t _(80)_ = −2.969*, *p = 0.0109,* Tukey correction method) (Figure 1B).

**Figure 1.**
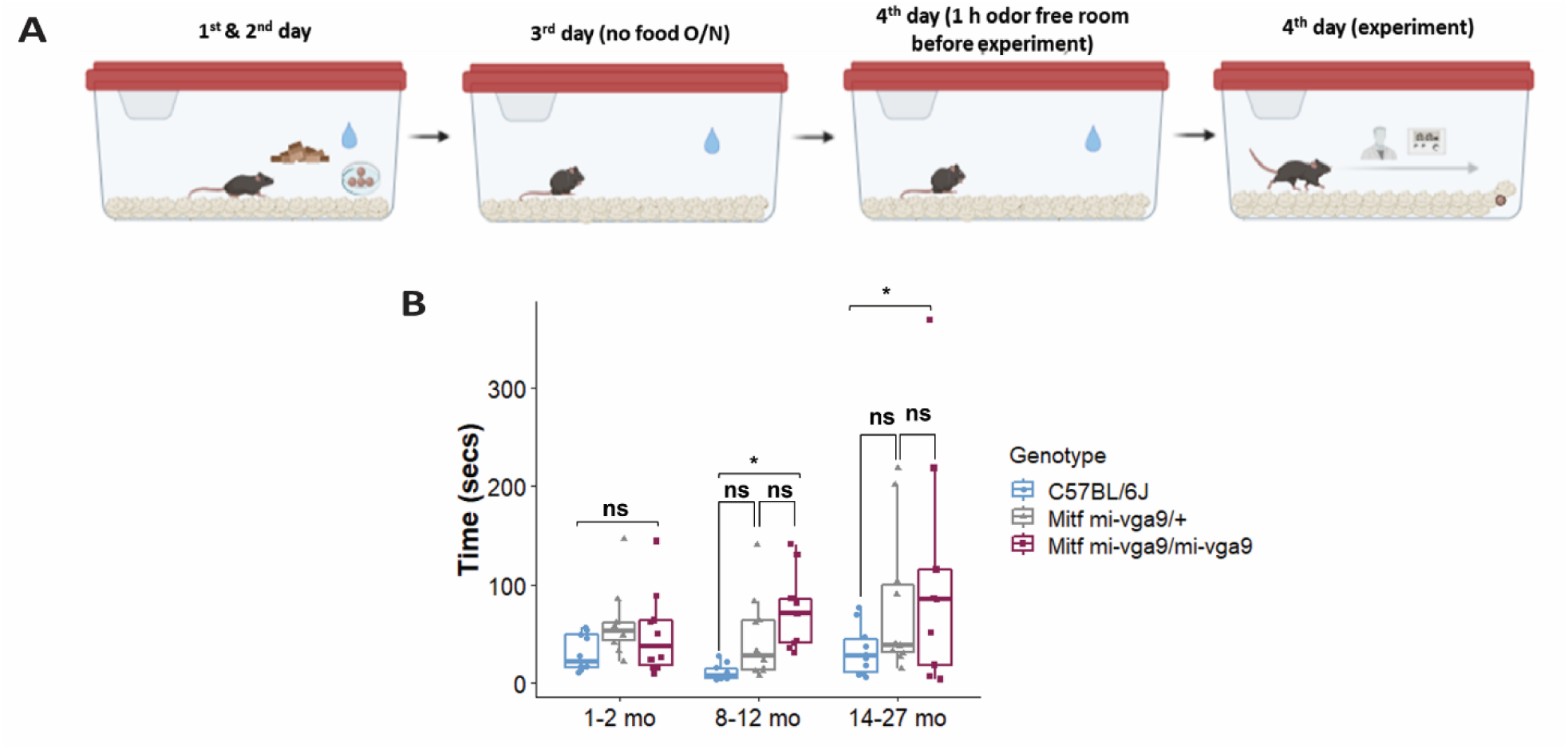
Aged *Mitf^mi-vga9/mi-vga9^* has reduced olfactory ability. A. Figure showing the setup of the hidden food assay. B. Results of the hidden food assay showing the time spent by three age groups of C57BL/6J, *Mitf^mi-vga9/+^* and *Mitf^mi-vga9/mi-vga9^* mice to find the hidden cereal. N=10 per genotype in age groups 1-2 mo and 8-12 mo; n=9 for 14-27 mo *Mitf^mi-vga9/mi-vga9^*. P-values were calculated using two-way ANOVA and adjusted with Tukey correction method. *p < 0.05, ns (not significant), mo (months old).

Variation in the time that *Mitf^mi-vga9/+^* and *Mitf^mi-vga9/mi-vga9^*mutant mice spent searching was high, some showing similar olfactory ability to C57BL/6J. In the aged groups there were individual *Mitf^mi-vga9/mi-vga9^* mice which showed severely reduced olfactory ability. Interestingly, *Mitf^mi-vga9/+^* mice, which are phenotypically similar to C57BL/6J (i.e., black coat color, normal eyes, melanocytes and mast cells presence), showed a trend of reduced olfactory ability. This suggests that the effect is primary, rather than secondary due to *Mitf* pleiotropy, and indicates a dosage effect associated with the *Mitf* mutation, which is in accordance with previous studies (Atacho et al., 2020; Ingason et al., 2019; Sabaté San José & Petersen, 2024).

### Potassium channel expression in aged *Mitf^mi-vga9/mi-vga9^* OB

Loss of *Mitf* leads to increased neuronal activity in primary PNs of young *Mitf^mi-vga9/mi-vga9^* OBs due to reduced I_A_ potassium currents (Atacho et al., 2020). Single molecule fluorescence in situ hybridization (smFISH) revealed decreased expression of potassium channel subunit *Kcnd3* (Kv4.3), correlating with increased neuronal activity in primary PNs of young *Mitf^mi-vga9/mi-vga9^*. On the other hand, expression of *Kcnd2* (Kv4.2) was increased in M/T cells of *Mitf^mi-vga9/mi-vga9^* mice, possibly due to a compensatory mechanism (Atacho et al., 2020). Given the hyperactivity in young *Mitf^mi-vga9/mi-vga9^*OB primary M/T cells (Atacho et al., 2020), global gene expression analysis of aged *Mitf^mi-vga9/mi-vga9^* and C57BL/6J OBs was performed to examine the effects of aging on the expression of potassium channel subunits (Figure 2A). Indications of inflammation or degeneration were also of interest.

**Figure 2.**
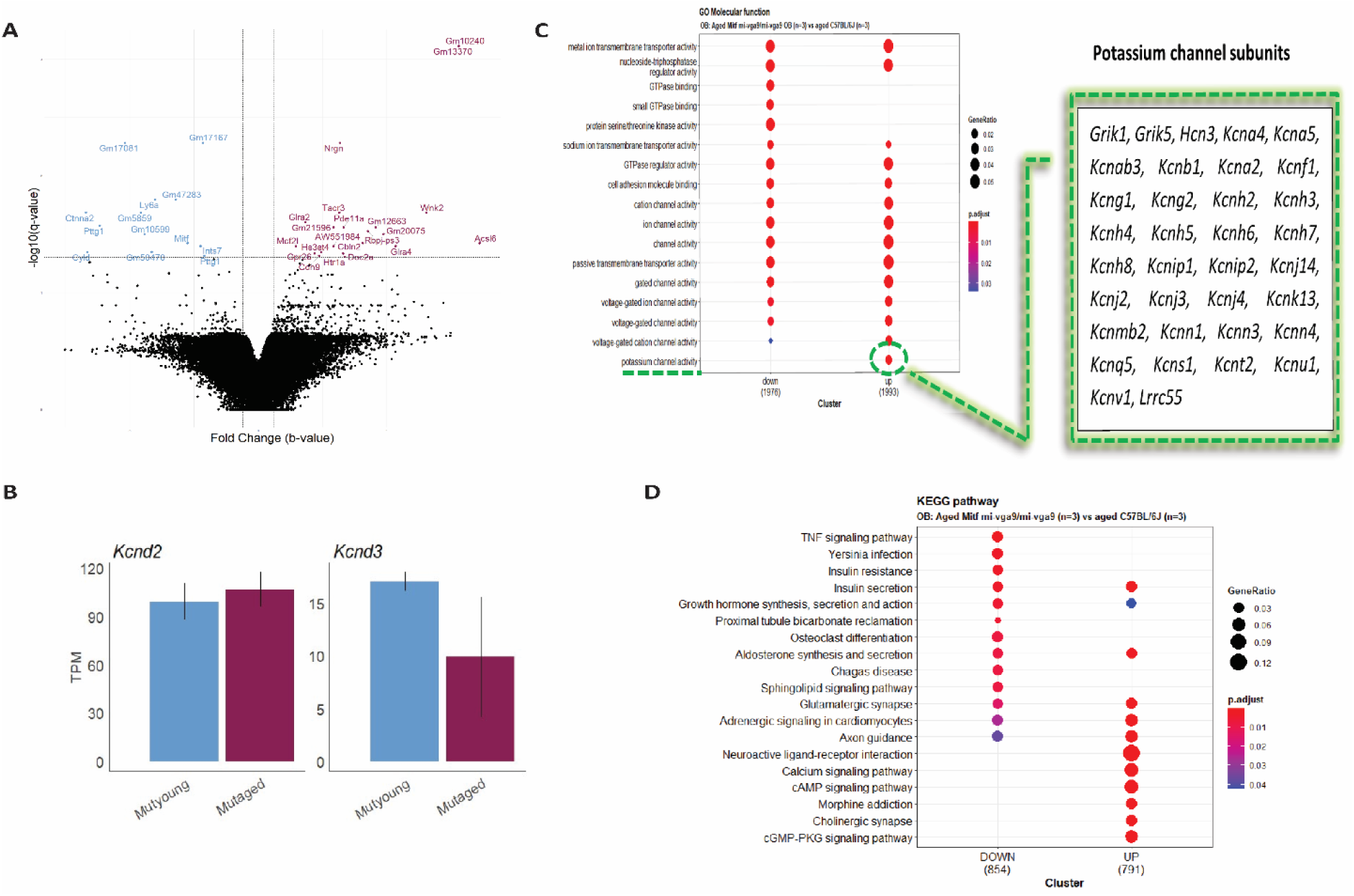
Increased potassium channel activity in aged *Mitf^mi-vga9/mi-vga9^* OB. A. Volcano plot shows DE genes in the OB of aged *Mitf^mi-vga9/mi-vga9^* compared to aged C57BL/6J. Blue indicates decreased genes, and purple indicates increased genes. N=3 per genotype. B. RNA expression of *Kcnd3* and *Kcnd2* in TPM in young (MUTyoung) and aged (MUTaged) *Mitf^mi-vga9/mi-vga9^*. N=3 per age group. C. GO Molecular Function (MF) analysis between aged *Mitf^mi-vga9/mi-vga9^* and aged C57BL/6J. Table shows genes enriched in the potassium channel activity cluster. D. KEGG pathway analysis between aged *Mitf^mi-vga9/mi-vga9^* and aged C57BL/6J. N=3 per genotype.

The potassium channel subunit (*Kcnd3*) previously shown to be reduced in expression in young *Mitf^mi-vga9/mi-vga9^* mice were not differentially expressed in aged *Mitf^mi-vga9/mi-vga9^* mice compared to aged C57BL/6J OB animals Their expression also remained unchanged when comparing aged and young *Mitf^mi-vga9/mi-vga9^* mice, suggesting that aging has no impact on global *Kcnd3* and *Kcnd2* expression (Figure 2B). A likely explanation is that the sensitivity and spatial resolution of smFISH allow this method to detect subtle changes in a specific cell types, which the global RNA-seq misses.

To investigate the biological functions of genes differentially expressed between aged *Mitf^mi-vga9/mi-vga9^* and C57BL/6J OBs, gene ontology (GO) and Kyoto Encyclopedia of Genes and Genomes (KEGG) pathway analysis were performed. GO molecular function (GO MF), identified overlapping categories between increased and decreased DE genes; the decreased DE genes were enriched in GTPase binding, small GTPase binding, and protein serine/threonine kinase activity. However, in general the expression of potassium channel subunits was increased in the aged *Mitf^mi-vga9/mi-vga9^*mice (Figure 2C). KEGG pathway analysis revealed that various pathways were enriched and again there was considerable significant overlap between the up- and downregulated DE genes. Pathways such as TNF signaling, Yersinia infection, insulin resistance, proximal tubule bicarbonate reclamation, osteoclast differentiation, Chagas disease, and the sphingolipid signaling pathway were associated with the decreased genes. Pathways such as neuroactive ligand-receptor interaction, calcium signaling pathway, morphine addiction, cholinergic synapse, and the cGMP-PKG signaling pathway were enriched by the increased DE genes (Figure 2D). However, no clear functional connection was observed with neurodegeneration or inflammation.

### Dendritic morphology in OBs does not change upon aging of *Mitf^mi-vga9/mi-vga9^* mice

Dendritic morphology can be affected by long term hyperactivity (Ugarte et al., 2023). To investigate this the Golgi-Cox method was used to evaluate the dendritic architecture of M/T cells in aged C57BL/6J and *Mitf^mi-vga9/mi-vga9^*OBs (Figure 3A) (Vints et al., 2019; Zhong et al., 2019). Several parameters were quantified, including the number of segments (Figure 3B), the segment length (Figure 3C), the number of branch points (Figure 3D), and tortuosity (Figure 3E) in each genotype. Analysis of these parameters revealed no differences between aged *Mitf^mi-vga9/mi-vga9^*and C57BL/6J OBs. Overall, the structure of dendrites was similar in both genotypes.

**Figure 3.**
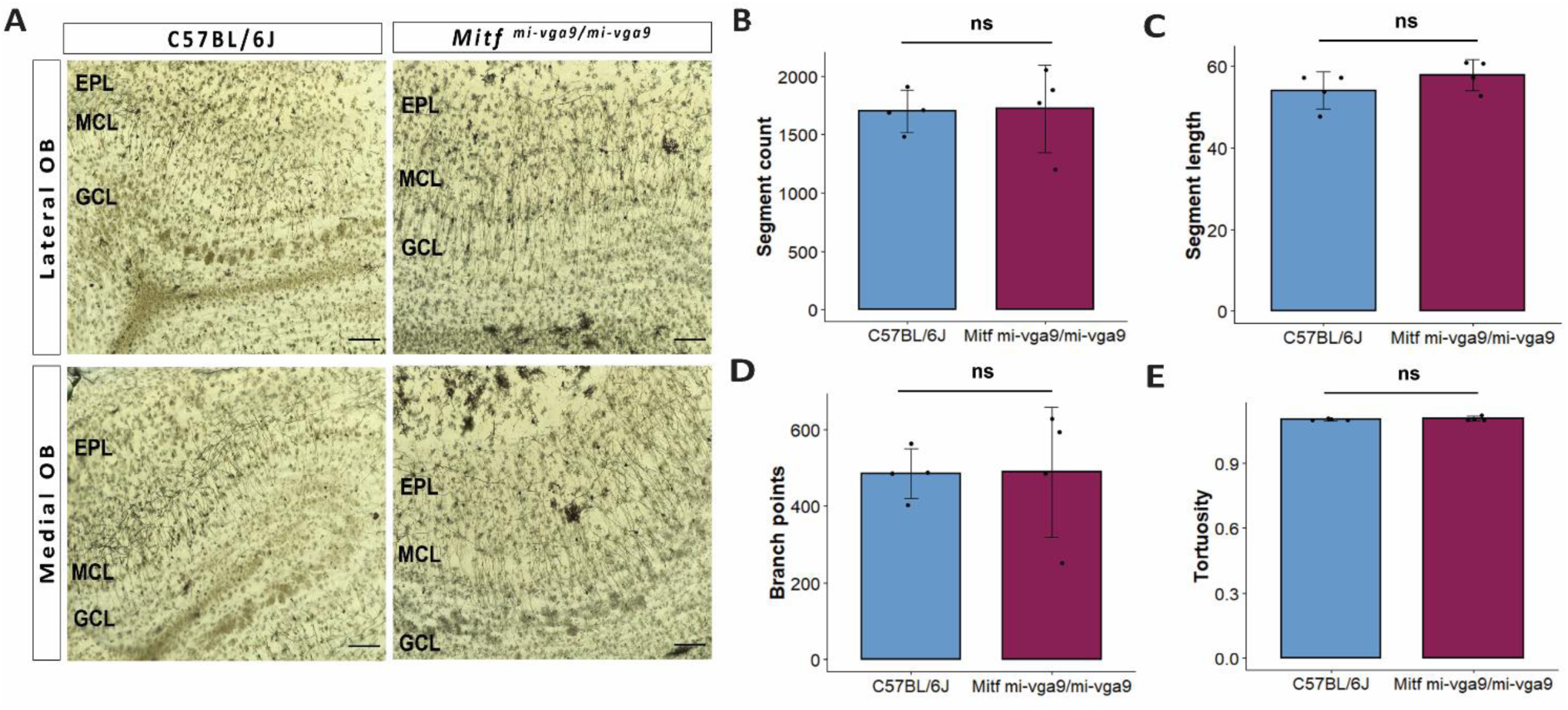
Dendritic morphology does not change upon aging of *Mitf^mi-vga9/mi-vga9^* OBs. A. Representative Golgi-Cox images from the lateral and medial OBs of aged C57BL/6J and *Mitf^mi-vga9/mi-vga9^* mice. Analysis of Golgi-Cox images with regards to the number of segments (B), segment length (C), number of branch points (D), and tortuosity (E). Scale bars: 200 μm. N=4 per genotype. Statistical analysis was performed using two-tailed unpaired Student’s t-test. ns (not significant).

### Inflammatory status does not change in *Mitf^mi-vga9/mi-vga9^* OBs upon aging

*Mitf* has been shown to be an important regulator of the neurodegenerative and phagocytic transcriptional program of microglia (Dolan et al., 2023). Long-term hyperactivity in neurons is a frequent first step in neurodegeneration (Targa Dias Anastacio et al., 2022). Given the involvement of MITF protein in the regulation of neuronal activity (Atacho et al., 2020), it is possible that the *Mitf* mutation may lead to neurodegeneration and subsequent inflammation, thereby contributing to the observed decline in olfactory ability.

To analyze the potential effects of *Mitf* on neurodegeneration and inflammation during aging, a comparative gene expression analysis using RNA-seq was performed to determine if there were any changes in genes associated to inflammation between aged C57BL/6J and *Mitf^mi-vga9/mi-vga9^*OBs. The analysis showed that there were no changes in the expression genes known to be specific to activated/reactive astrocytes (Figure 4A) or microglia activation (Figure 4B) between aged *Mitf^mi-vga9/mi-vga9^*and C57BL/6J OBs. This confirmed that the inflammatory status of *Mitf^mi-vga9/mi-vga9^* OBs remains unchanged upon aging.

**Figure 4.**
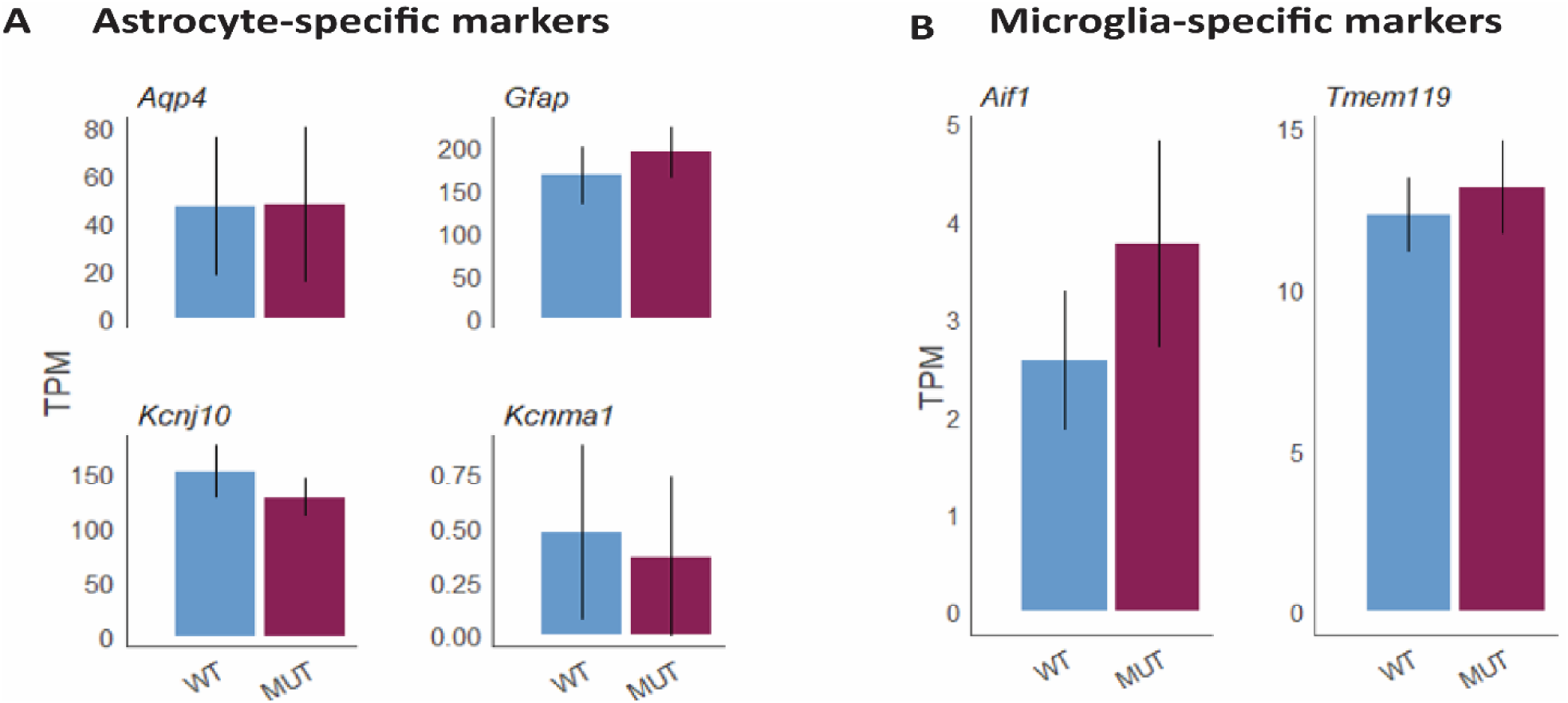
No change in gene expression level of markers for astrocytes and microglia at the gene level in aged *Mitf^mi-vga9/mi-vga9^* OB. Expression of astrocyte-specific markers (*Aqp4, Gfap, Kcnj10, Kcnma1*) (A) and microglia-specific markers (*Aif1* aka *Iba-1*, *Tmem119*) (B) in TPMs in aged *Mitf^mi-vga9/mi-vga9^*(MUT) OB compared to C57BL/6J (WT) OB. N=3 per genotype.

### No evidence of neuroinflammation in aged *Mitf^mi-vga9/mi-vga9^* OB

To further examine inflammation at the protein level, immunohistochemistry was performed on OB sections of young and aged C57BL/6J and *Mitf^mi-vga9/mi-vga9^* using well-established neuroinflammation markers. Neuroinflammation and neurodegeneration in the CNS are generally studied by analyzing the state of astrocytes and microglia (Lucas et al., 2006) using GFAP, a marker for astrocytes and astrogliosis (Rodríguez et al., 2014), and Iba-1, a marker used to identify and characterize microglia (Streit et al., 2009). No changes were observed in the number of astrocytes in the OBs of young (Figure 5A) and aged (Figure 5B) C57BL/6J or *Mitf^mi-vga9/mi-vga9^* mice. These results were confirmed with western blot analysis of GFAP, which showed no changes in expression in young (Figure 5C, E) or aged (Figure 5D, F) *Mitf^mi-vga9/mi-vga9^* and C57BL/6J OBs. Likewise, there were no changes in the number of microglia between OBs of young (Figure 6A) or aged (Figure 6B) *Mitf^mi-vga9/mi-vga9^* or C57BL/6J mice. Similar results were obtained in the western blot analysis of Iba-1. No significant difference was observed in the expression of Iba-1 between young (Figure 6C, E) and aged (Figure 6D, F) *Mitf^mi-vga9/mi-vga9^*and C57BL/6J OBs. In summary, there was no evidence of neuroinflammation in aged *Mitf^mi-vga9/mi-vga9^* OBs based on the expression of glia markers.

**Figure 5.**
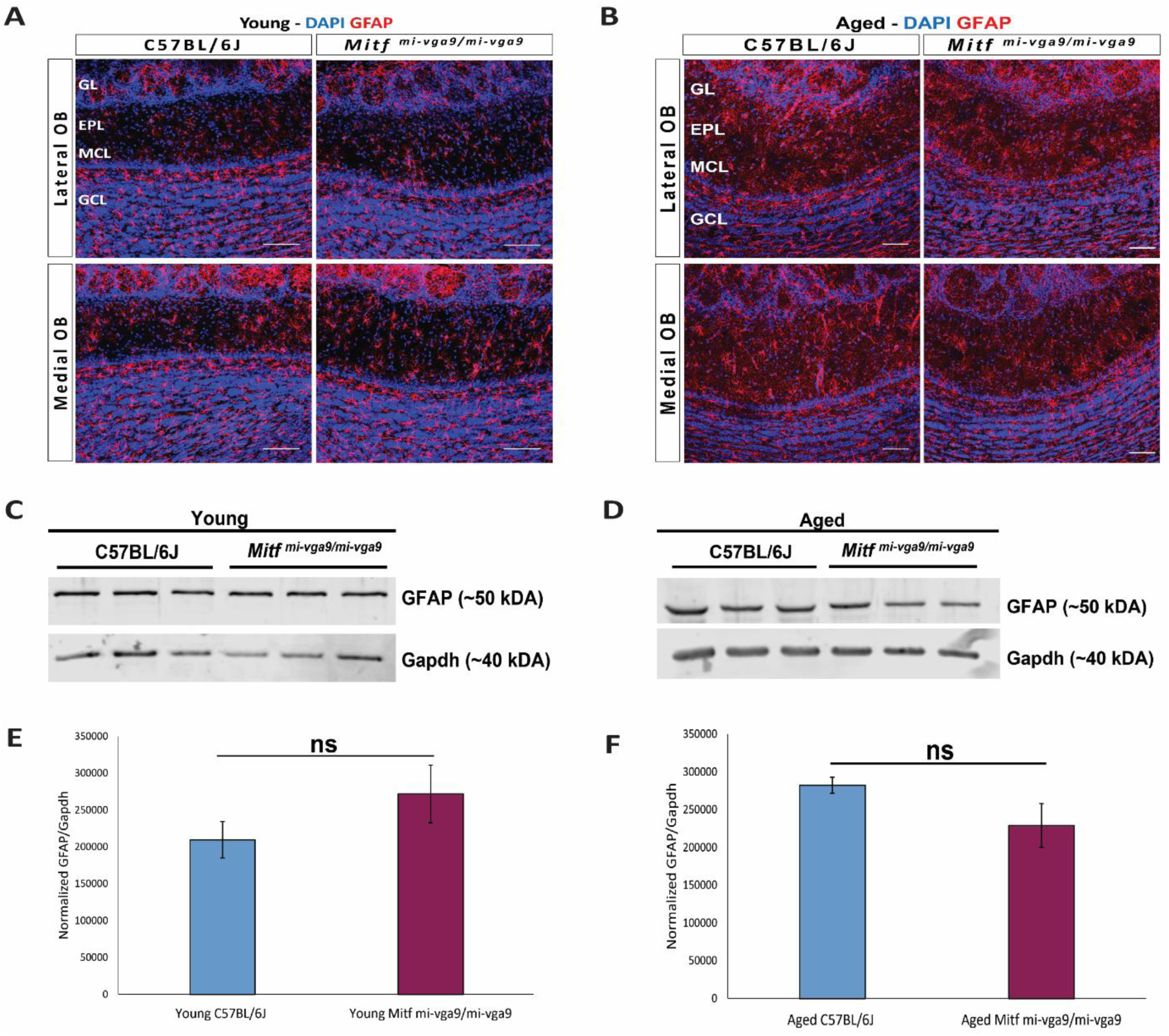
Expression of the astrocyte/astrogliosis marker is unchanged in young and aged *Mitf^mi-vga9/mi-vga9^* OB. Representative immunofluorescence images show the expression of GFAP (red) protein in young (A) and aged (B) C57BL/6J and *Mitf^mi-vga9/mi-vga9^* OB. N=10 per genotype and age group. DAPI nuclear staining is shown in blue. Scale bars: 100 μm. Representative images of immunoblots from whole lysate of young (C) and aged (D) C57BL/6J and *Mitf^mi-vga9/mi-vga9^* OB probed for GFAP and Gapdh as a loading control. Densitometric analysis of GFAP bands normalized to the Gapdh loading control in young (E) and aged (F) C57BL/6J and *Mitf^mi-vga9/mi-vga9^*. All immunoblotting experiments were performed with three mice (n=3) per genotype and age group. Statistical analysis was performed using two-tailed unpaired Student’s t-test. ns (not significant).

**Figure 6.**
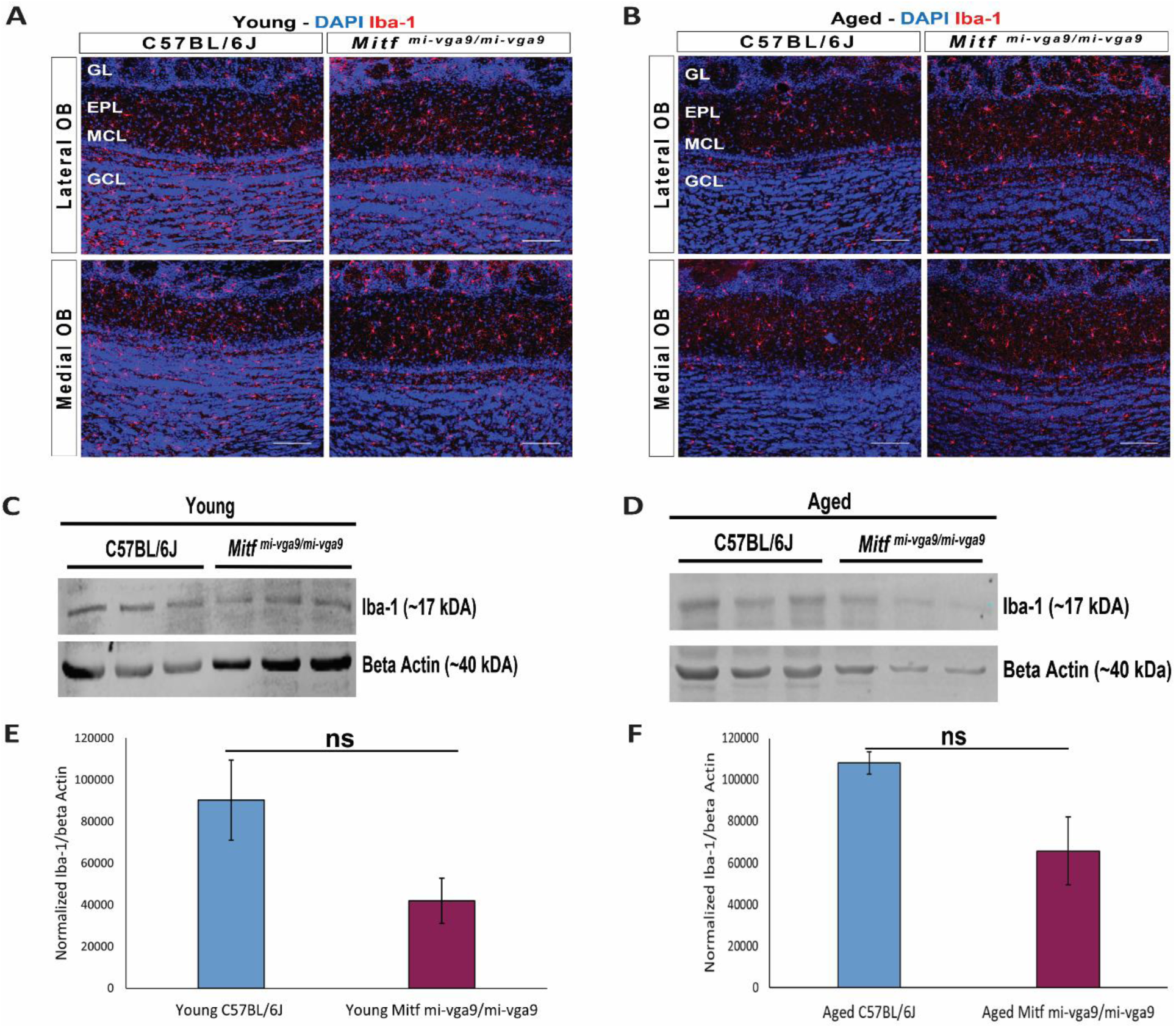
Expression of the microglia activation marker (Iba-1) marker does not change in young and aged *Mitf^mi-vga9/mi-vga9^* OBs. Representative immunofluorescence images for Iba-1 (red) protein in young (A) and aged (B) C57BL/6J and *Mitf^mi-vga9/mi-vga9^* OB. N=10 per genotype and age group. DAPI nuclear staining is shown in blue. Scale bars: 100 μm. Representative images of immunoblotting from whole lysate of young (C) and aged (D) C67BL/6J and *Mitf^mi-vga9/mi-vga9^* OB probed for Iba-1 and beta-Actin as a loading control. Densitometric analysis of Iba-1 bands normalized to beta-Actin in young (E) and aged (F) C57BL/6J and *Mitf^mi-vga9/mi-vga9^*OBs. All immunoblotting experiments were performed with three mice (n=3) per genotype and age group. Statistical analysis was performed using two-tailed unpaired Student’s t-test. ns (not significant).

### Quantification of PNs in aging OB

Neurodegeneration can result in loss of neurons (Yankner et al., 2008). To investigate this in aged mice, OB tissues from young and aged C57BL/6J and *Mitf^mi-vga9/mi-vga9^* mice were stained with antibody against the Tbr2/Eomes protein, a marker for OB PNs (Figure 7A). Prior findings demonstrated an increase in Tbr2/Eomes positive cells in the GL but not in the EPL or MCL of young *Mitf^mi-vga9/mi-vga9^* OBs (Atacho et al., 2020). However, the number of Tbr2/Eomes positive cells was not changed in GL, EPL, or MCL of aged *Mitf^mi-vga9/mi-vga9^*when compared to aged C57BL/6J OBs (Figure 7B).

**Figure 7.**
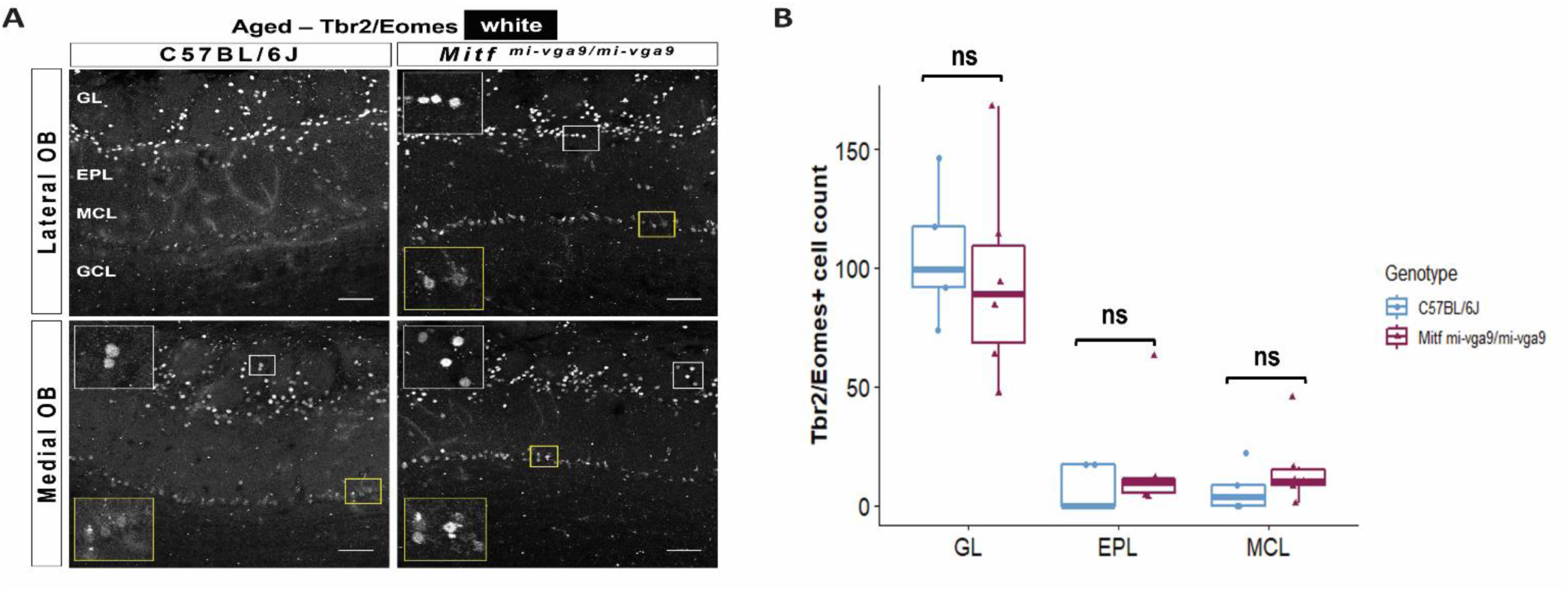
The number of Tbr2/Eomes positive cells does not change upon aging. A. Representative immunofluorescence images for Tbr2/Eomes (white) protein in aged C57BL/6J and *Mitf^mi-vga9/mi-vga9^* OBs. B. Quantification of Tbr2/Eomes positive cells in the GL, EPL, MCL of aged C57BL/6J and *Mitf^mi-vga9/mi-vga9^* OBs. N=5 for C57BL/6J and n=6 for *Mitf^mi-vga9/mi-vga9^*. Scale bars: 100 μm. Statistical analysis was performed using two-way ANOVA. ns (not significant).

The OBs from young (Figure 8A) and aged (Figure 8B) C57BL/6J and *Mitf^mi-vga9/mi-vga9^* mice were stained with Tbr1, a marker of OB PNs. This staining showed no changes in the number of Tbr1 positive cells in the GL, EPL, or MCL between young *Mitf^mi-vga9/mi-vga9^* and C57BL/6J OBs (Figure 8C). However, the number of Tbr1 positive cells was significantly increased in the GL in aged *Mitf^mi-vga9/mi-vga9^*OBs, whereas no change was observed in the EPL and MCL (Figure 8D; *t _(12)_ = −3.987*, *p = 0.0267*, Sidak multiple comparison). Taken together, there was no reduction in the number of PNs, arguing against neuronal degeneration.

**Figure 8.**
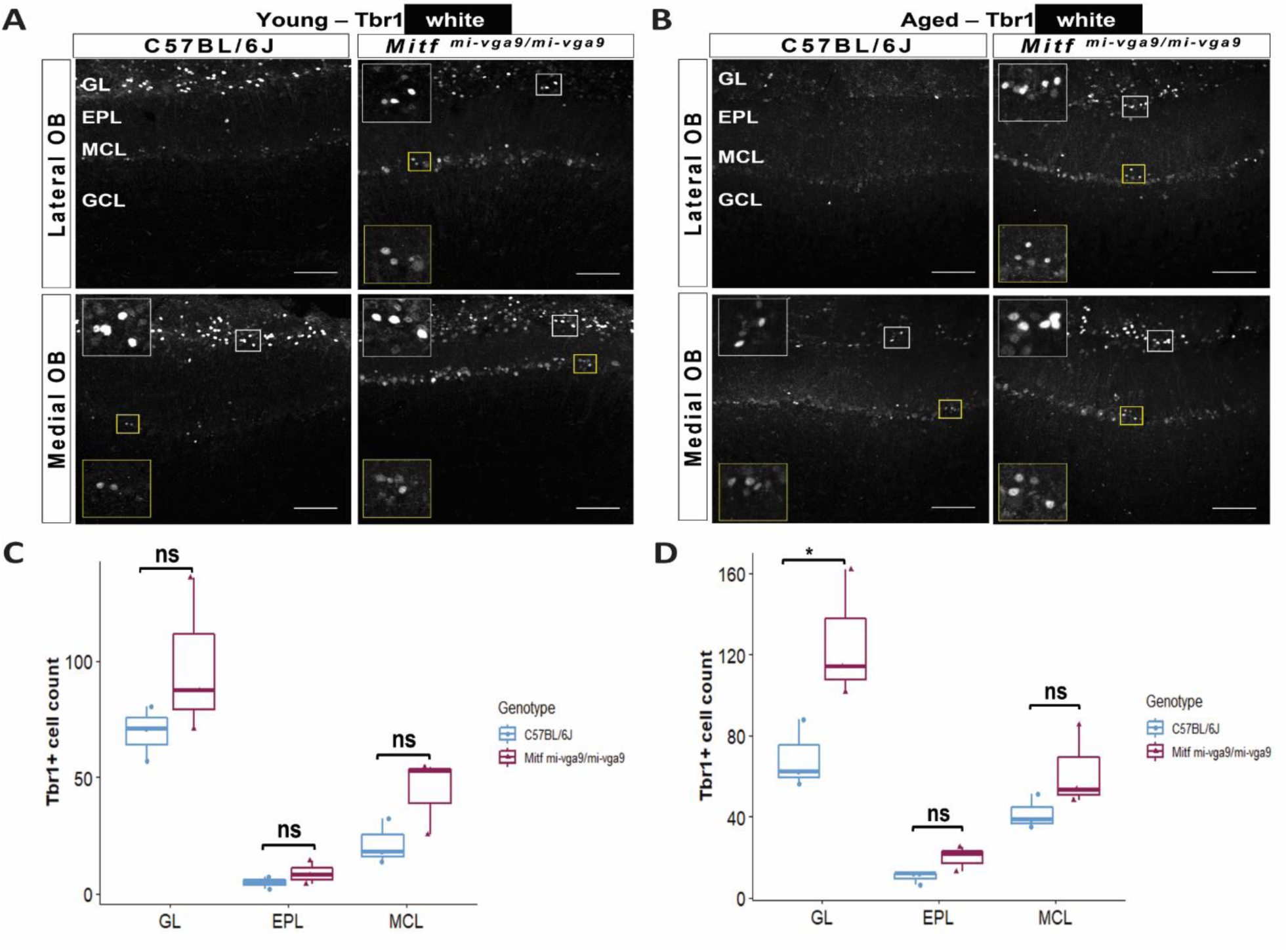
The number of Tbr1 positive cells is increased in the GL of aged *Mitf^mi-vga9/mi-vga9^* OBs. Representative immunofluorescence images for Tbr1 (white) protein on young (A) and aged (B) C57BL/6J and *Mitf^mi-vga9/mi-vga9^* OBs. Quantification of Tbr1 positive cells in the GL, EPL, MCL of young (C) aged (D) C57BL/6J and *Mitf^mi-vga9/mi-vga9^* OBs. N=3 per genotype and age group. Scale bars: 100 μm. P-values were calculated using two-way ANOVA and adjusted with Sidak multiple comparison. *p < 0.05, ns (not significant).

## Discussion

Reduction in olfactory ability, neuroinflammation, and neuronal hyperactivity often follows or precedes neurodegeneration, e.g., in AD (Doty, 2012; Fullard et al., 2017; Murphy, 2019; Patino et al., 2021; Peters et al., 2003). Aging *Mitf^mi-vga9/mi-vga9^* mice often have reduced olfactory ability and primary PNs from young *Mitf^mi-vga9/mi-vga9^* OB are hyperactive (Atacho et al., 2020). This could lead to increased neuronal or neural circuit stress *in vivo* and subsequently to neurodegeneration. The effects of *Mitf* are pleiotropic and loss of *Mitf* could also lead to degeneration through other mechanisms (Dolan et al., 2023) resulting in the reduction of olfactory ability.

The current study shows that aged *Mitf^mi-vga9/mi-vga9^*OBs have reduced olfactory ability with no clear evidence of neuroinflammation or neurodegeneration. No increase was detected in inflammation upon aging of *Mitf^mi-vga9/mi-vga9^* mice, neither at the gene level nor in expression of inflammatory proteins. Similarly, no reduction was observed in the number of OB PNs in *Mitf^mi-vga9/mi-vga9^*mice with aging. Comparing gene expression between aged C57BL/6J and aged *Mitf^mi-vga9/mi-vga9^* mice showed a high number of DE genes, with diverse functions and connected to diverse pathways. However, as the loss of *Mitf* leads to changes in the expression of two genes coding for potassium channel subunits in M/T cells, it was particularly interesting that there was a global increase in the expression of genes coding for potassium channel subunits during aging. Overall, such a change should lead to reduced neuronal activity. This suggests that with aging, neuronal hyperactivity leads to a reduction in neuronal activity. As complex mechanisms regulate the conduction of information in the OB, these changes are likely to be multifaceted (Burton, 2017) involving complicated interactions between various inhibitory neurons and neurotransmitter systems, contributing to the delicate balance of excitation and inhibition necessary for proper sensory processing (Abraham et al., 2010; Egger et al., 2003; Geramita & Urban, 2017). The disruption of this equilibrium could underlie the observed decline in olfactory function (Gire et al., 2019) in aged *Mitf^mi-vga9/mi-vga9^* mice. Other possibilities could be a decline in the function of the OE or OSNs. The *Mitf* mutation may also elicit effects potentially impacting processes such as proteostasis. MITF has been associated with autophagy in other cell types, including in melanoma cells (Möller et al., 2019). However, no specific changes were observed in the expression of genes that may explain such effects.

A caveat to this study is secondary effects due to the pleiotropy of *Mitf*. That the olfactory phenotype is also present in the heterozygous *Mitf ^mi-vga9^* animals, phenotypically similar to C57BL/6J mice, supports this to be a primary effect. It can be concluded from the current study that the absence of *Mitf* does not lead to neurodegeneration with aging. However, *Mitf* could play a role in neurodegeneration induced by neuropathology, as in AD (Dolan et al., 2023). A direct challenge in a conditional model may unravel this.

## Acknowledgements

This study was supported in part by Research Fund of Iceland to PHP (217945-052) and a grant from the University of Iceland Doctoral Grants Fund to FM. We thank the Biomedical Center at the University of Iceland (BMC) and Utopia Arctica for their contribution to data analysis. We are also grateful to our colleagues from the BMC Ragnhildur Káradóttir, Margrét Helga Ögmundsdóttir, Sigríður Rut Franzdóttir, and Sana Gadiwalla for their insights and assistance during the project.

